# SimpleSBML: A Python package for creating and editing SBML models

**DOI:** 10.1101/030312

**Authors:** Caroline Cannistra, Kyle Medley, Herbert M. Sauro

## Abstract

**Summary:** In this technical report we describe a simple extension to python-libSBML that allows users of Python to more easily construct SBML based models. The most commonly used package for constructing SBML models in Python is python-libSBML based on the C/C++ library libSBML. python-libSBML supports a comprehensive set of model types, but is difficult for new users to learn and requires long scripts to create even the simplest models. We present SimpleSBML, a package that allows users to add species, parameters, reactions, events, and rules to a libSBML model with only one command for each. Models can be exported to SBML format, and SBML files can be imported and converted to SimpleSBML commands that creates each element in a new model. This allows users to create new models and edit existing models for use with other software.

**Accessibility and Implementation:** SimpleSBML is publicly available and licensed under the liberal Apache 2.0 open source license. It supports SBML levels 2 and 3. Its only dependency is libSBML. It is supported on Windows and Mac OS X. All code has been deposited at the GitHub site https://github.com/sys-bio/simplesbml

## 1 Introduction

Biological modeling is a key component of systems biology, and the development of Systems Biology Markup Language (SBML) [1], a markup language designed to describe models of biological systems, has allowed systems and synthetic biologists to develop a plethora of useful software tools that are automatically compatible with each other through SBML document import and export. Creating SBML models is possible with a variety of software packages, the most well-known of which is libSBML [2]. This package allows users to generate SBML documents by writing scripts in Python,C or C++. While libSBML is useful, if can be difficult to learn for novices and even the simplest of models using libSBML tend to be very long. SimpleSBML is a package that also allows users to create SBML models with Python scripting, but requires far fewer commands and is more accessible for beginner users.

SimpleSBML makes use of libSBML methods to add elements such as species, parameters, reaction, events and rules to a model, and generate an SBML-formatted document from the resulting model object. SimpleSBML also includes a method that imports an SBML document and generates SimpleSBML code that corresponds to the model described in the document. This allows users to edit SBML-formatted models as well as create new ones.

## 2 Methods

### 2.1 sbmlModel methods

Users can create new models with the sbmlModel class, which holds a SBMLDocument object and contains methods that add different elements, such as species, parameters, reactions and events.

Here is an example of a simple reaction-based model built with sbmlModel.

~~~
import simplesbml
model = simplesbml.sbmlModel ();
model.addCompartment(1e-14, comp_id=’comp’);
model.addSpecies(’E’, 5e-21, comp=’comp’);
model.addSpecies(’S’, 1e-20, comp=’comp’);
model.addSpecies(’P’, 0.0, comp= ’ comp’);
model.addSpecies(’ES’, 0.0, comp=’comp’);
model.addReaction([’E’, ’S’], [’ES’], ’comp*(kon*E*S-koff*ES)’, \
              local_params= {’koff’: 0.2, ’kon’: 1000000.0}, rxn_id=’veq’);
model.add Reaction([’ES’], [’E’, ’P’], ’comp*kcat*ES’, \
                           local_params= {’kcat’: 0.1}, rxn_id=’vcat’);
~~~

In this example, reaction rate constants are stored locally with the reactions where they are used. It is also possible to define global parameters and use them in reaction expressions. Here is an example of this usage.

~~~
import simplesbml
model = simplesbml.sbmlModel ();
model.addCompartment(1e-14, comp_id=’comp’);
model.addSpecies(’E’, 5e-21, comp=’comp’);
model.addSpecies(’S’, 1e-20, comp=’comp’);
model.addSpecies(’P’, 0.0, comp= ’ comp’);
model.addSpecies(’ES’, 0.0, comp=’comp’);
model.addParameter(’koff’, 0.2);
model.addParameter(’kon’, 1000000.0);
model.addParameter(’kcat’, 0.1);
model.addReaction([’E’, ’S’], [’ES’], ’comp*(kon*E*S-koff*ES)’, rxn_id=’veq’);
model.addReaction([’ES’], [’E’, ’P’], ’comp*kcat*ES’, rxn_id=’vcat’);
~~~

SimpleSBML also supports the use of events to change the system state under certain conditions, and the use of assignment rules and rate rules to explicitly define variable values as a function of the system state. Here is an example of events and rate rules. In this example, the value of parameter G2 is instantaneously determined by the relationship between P1 and tau, and the rates of change of P1 and P2 are explicitly defined in equation form instead of using a reaction.

~~~
import simplesbml
model = simplesbml.sbmlModel ();
model.addCompartment (vol=1.0, comp_id=’cell’);
model.addSpecies(’[P1]’, 0.0, comp=’cell ’);
model.addSpecies(’[P2]’, 0.0, comp=’cell ’);
model.addParameter(’k1’, 1.0);
model.addParameter(’k2’, 1.0);
model.addParameter(’tau’, 0.25, units=’concentration’);
model.addParameter(’G1’, 1.0, units=’concentration’);
model.addParameter(’G2’, 0.0, units=’concentration’);
model.addEvent(trigger=’P1 > tau’, assignments={’G2’: ’1 concentration’});
model.addEvent(trigger=’P1 <= tau’, assignments={’G2’: ’0 concentration’});
model.addRateRule(’P1 ’, ’k1 * (G1 - P1)’);
model.addRateRule(’P2’, ’k2 * (G2 - P2)’);
~~~

The following example shows how it is possible to easily encode a standard differential equation model, in this case the Lorenz model (https://en.wikipedia.org/wiki/Lorenz_system) using SimpleSBML. Once described in this way the SBML model can be loaded into a Python based simulator such as libRoadRunner [3]. Note the use of toSBML to retrieve the SBML string.

~~~
import simplesbml
model = simplesbml.sbmlModel ();
model.addSpecies(’x’, 0.96259);
model.addSpecies(’y’, 2.07272);
model.addSpecies(’z’, 18.65888);
model.addParameter(’sigma’, 20.0);
model.addParameter(’rho’, 28);
model.addParameter(’beta’, 2.67);
model.addRateRule(’x’, ’sigma*(y - x)’);
model.addRateRule(’x’, ’x*(rho - z) - y’);
model.addRateRule(’x’, ’x*y - beta*z’);
sbmlStr = model.toSBML ()
~~~

### 2.2 writeCode

Users can edit existing models with the writeCode() method, which accepts a SBML document and produces a script of SimpleSBML commands in string format. This method converts the SBML document into a libSBML Model and scans through its elements, adding lines of code for each SimpleSBML-compatible element it finds. The output can be saved to a .py file and edited to create new models based on the original import. For instance, here is an example of a short script that reproduces the SimpleSBML code to reproduce an sbmlModel object.

~~~
import simplesbml
model = simplesbml.sbmlModel ();
model.addCompartment(1e-14, comp_id=’comp’);
model.addSpecies(’E’, 5e-21, comp=’comp’);
model.addSpecies(’S’, 1e-20, comp=’comp’);
model.addSpecies(’P’, 0.0, comp= ’ comp’); model.addSpecies(’ES’, 0.0, comp=’comp’);
model.addReaction([’E’, ’S’], [’ES’], ’comp*(kon*E*S-koff*ES)’, \
             local_params={’koff’: 0.2, ’kon’: 1000000.0}, rxn_id=’veq’);
model.addReaction([’ES’], [’E’, ’P’], ’comp*kcat*ES’, \
                          local_params={’kcat’: 0.1}, rxn_id=’vcat’);
code = simplesbml.writeCodeFromString(model.toSBML());
f = open(’example_code.py’, ’w’);
f.write(code);
f.close ();
The output saved to ’example_code.py’ will look like this:
import simplesbml
model = simplesbml.sbmlModel(sub_units=’’);
model.addCompartment(vol=1e-14, comp_id=’comp’);
model.addSpecies(species_id=’E’, amt=5e-21, comp=’comp’);
model.addSpecies(species_id=’S’, amt=1e-20, comp=’comp’);
model.addSpecies(species_id=’P’, amt=0.0, comp=’comp’);
model.addSpecies(species_id=’ES’, amt=0.0, comp=’comp’);
model.addReaction(reactants=[’E’, ’S’], products=[’ES’], \
             expression=’comp * (kon * E * S - koff * ES)’, \
             local_params={’koff’: 0.2, ’kon’: 1000000.0}, rxn_id=’veq’);
model.addReaction(reactants=[’ES’], products=[’E’, ’P’], \
             expression=’comp * kcat * ES’, local_params={’kcat’: 0.1}, \
             rxn_id=’vcat ’);
~~~

## 3 Installation

SimpleSBML depends on libSBML. In order to install SimpleSBML, libSBML for python must first be installed. Refer to the https://pypi.python.org/pypi/python-libsbml for details. SimpleS-BML can be downloaded as a .zip from https://github.com/sys-bio/simplesbml. To install unzip the contents, or directly clone the directory and copy the contents to a convenient place on your hard drive. At the command line, move into the ’simplesbml/’ folder. This folder should also contain a file called ’setup.py’. Type ’python setup.py install’. Make sure that Python is on the search path so that setup.py can be executed. To use simpleSBML import the module in Python by typing ’import simplesbml’ into the command line in Python. SimpleSBML is also available ready to use from the Tellurium modeling package found at http://tellurium.analogmachine.org/.

## 4 Discussion

SimpleSBML is intended for systems biology researchers who have limited experience with programming, or are working on simple models and prefer to use a simpler set of commands compared to libSBML. Future versions of SimpleSBML may include support for algebraic rules, conversion between different levels and versions of SBML, and future libSBML and SBML features as they are released.

## Supplementary information

Documentation and download is available at sys-bio.github.io/simplesbml.

## Acknowledgments

Funding: The authors are most grateful to generous funding from the National Institute of General Medical Sciences of the National Institutes of Health under award R01-GM081070. The content is solely the responsibility of the authors and does not necessarily represent the official views of the National Institutes of Health.

